# Climatic Influence on COVID-19: Investigating the Role of Temperature and Humidity in the Spread of the Omicron Variant

**DOI:** 10.1101/2025.09.11.675496

**Authors:** Leonardo López, Brandon Jersai García Checa, Xavier Rodó

**Affiliations:** Catalan Institution for Research and Advanced Studies (ICREA), Barcelona, Spain; Barcelona Institute of Global Health (ISGlobal), Barcelona, Spain; Barcelona University (UB), Barcelona, Spain

**Keywords:** Omicron, COVID-19, SARS-CoV-2, Climate, Temperature, Absolute Humidity, Model, Gobal

## Abstract

Understanding how climate modulates infectious disease dynamics is critical for anticipating epidemic patterns. This study examines the association between climatological variables—specifically temperature and relative humidity—and the incidence of the SARS-CoV-2 Omicron variant (*B*.1.1.529) during its global wave (2021–2022). Using global epidemiological and climate data, we applied Scale-Dependent Correlation (SDC) analysis to detect transient, scale-specific associations across regions and periods. We identified consistent negative correlations between incidence and both temperature and humidity, especially in mid-latitudes during colder months. These findings were compared with predictions from stochastic population-based compartmental models incorporating climate-dependent transmission parameters. Among the tested formulations, the temperature-based model achieved the best fit to observed case trajectories. Our results highlight a robust climatological influence on Omicron transmission dynamics and underscore the importance of integrating climate indicators into epidemic modeling and preparedness strategies.

## 1. Introduction

Pandemics have historically shaped human societies, economies, and public health strategies. The Black Death in the 14*th* century and the Spanish Flu of 1918 are stark reminders of the devastating impact infectious diseases can have on global populations (Piret and Boivin, 2021). More recently, the H1N1 swine flu pandemic of 2009 and especially the COVID-19 pandemic have further illustrated the profound societal and economic disruptions caused by such outbreaks (Sampath et al., 2021). These events, which claimed millions of lives, have also prompted significant advancements in epidemiology and public health infrastructure, underscoring the importance of preparedness and response strategies in mitigating the effects of future pandemics.

The relationship between climate variables and infectious disease dynamics has long been a subject of significant interest (Lam et al., 2020). Factors like temperature, humidity, radiation, and precipitation can influence the transmission and survival of pathogens in the environment Lam et al. (2020); Thomas (2020). For instance, colder temperatures and lower humidity have increased transmission rates of respiratory viruses such as influenza (Axelsen et al., 2014) . A crucial moment during the pandemic was the growing recognition of the airborne nature of the virus, which shifted the understanding of transmission mechanisms and public health responses (Wang et al., 2021).

The global community has recently encountered new challenges from emerging infectious diseases (Piret and Boivin, 2021). The rapid spread of pathogens across continents has been facilitated by international travel and urbanization (Sampath et al., 2021). The COVID-19 pandemic, caused by the novel coronavirus SARS-CoV-2, exemplifies how well interconnected our world is and highlights the deep impact a novel infectious agent can have on global health and economies (Ciotti et al., 2020; Pokhrel and Chhetri, 2021).

Recent research has identified climatic signatures linked to COVID-19 spread, revealing that colder temperatures and lower humidity are associated with increased transmission during earlier pandemic waves (Fontal et al., 2021). These findings determined critical temperature ranges (12-18°C) and absolute humidity levels (4 − 12*g/m*^3^) where transmission rates were notably heightened Fontal et al. (2021); Bashir et al. (2020); Briz-Redón and Serrano-Aroca (2020).

The emergence of genetic variants of SARS-CoV-2, such as those with the *D*614*G* mutation, marked a turning point in the pandemic’s trajectory (Tao et al., 2021). Variants of concern (VoCs) like Alpha, Beta, Gamma, Delta, and most recently, Omicron, have posed new challenges with their increased transmissibility, altered disease severity, and potential immune evasion capabilities (Choi and Smith, 2021).

The Omicron variant,first detected in Botswana and South Africa in November 2021, quickly spread globally, causing a large surge in cases (Karim and Karim, 2021). Epidemiological data indicated that Omicron exhibited significantly higher transmissibility than the Delta variant (Antonelli et al., 2022). Omicron’s estimated basic reproduction number (*R*_0_) ranges from 9.5 to 10.0, whereas Delta’s *R*_0_ was around 5.08. The consensus average effective reproduction number (*R*_*e*_) for Omicron ranges between 3.4 and 4.2, significantly higher than Delta’s *R*_*e*_ of about 1.25 − 1.57 (Antonelli et al., 2022; Karim and Karim, 2021). This heightened transmissibility prompted the World Health Organization to designate Omicron as a Variant of Concern on November 26, 2021.

Figure S1 illustrates the phylogenetic relationships among various SARS-CoV-2 variants, highlighting the evolutionary paths and connections between different lineages. The Omicron variant (*B*.1.1.529), marked in yellow, stands out with many sub-lineages, indicating its significant mutation rate and adaptability. The original Omicron variant and its subsequent sub-lineages have become dominant strains in various regions worldwide. Understanding these evolutionary relationships and the potential impact of climate on the dynamics of the Omicron strain appears crucial, as it helps underscore the role of environmental factors in the spread and evolution of this highly transmissible variant.

While previous research identified climatic signatures associated with COVID-19 spread during earlier pandemic waves, these studies could not account for the emergence of new variants (Fontal et al., 2021). With a significantly higher degree of transmissibility, the Omicron variant presents a unique challenge in understanding how climate variables influence its population propagation. In this sense, this study aims to investigate whether the climate-virus relationships observed earlier also persisted or differed during the Omicron wave. By examining these interactions on a global scale, we aim to provide relevant insights that can inform targeted public health interventions and policy decisions, ultimately aiding in the combat of new COVID-19 strains while preparing for future pandemics involving respiratory viruses.

Early studies at the beginning of the pandemic found no significant relationships between temperature and virus spread in China (Shi et al., 2020; Sera et al., 2021). However, more recent literature supported significant associations between SARS-CoV-2 transmission and weather conditions, particularly temperature and humidity (Fontal et al., 2021; Sera et al., 2021). High temperature and humidity were shown to negatively affect virus transmission, while high temperature alone also appeared to produced similar effects on viral propagation (Fontal et al., 2021). Instead, decreases in both absolute humidity and temperatures in midlatitudes were seen to be associated with viral spread irrespective of populations and countries, therefore providing support to the role of physical conditions sustaining longer times aerosol particles emitted by infectious patients and favoring short-term contagion (Morawska and Milton, 2020; Morawska et al., 2020).

Research on the role of climate in COVID-19 transmission dynamics, particularly in African countries, remains under-documented. Some studies, albeit controversy continues (Koanda et al., 2023; Mwiinde et al., 2022; Limaheluw et al., 2024), suggest significant albeit complex associations between meteorological factors and COVID-19 transmission across Africa, demonstrating the not fully resolved interplay between climate variables and COVID-19 dynamics.

As mentioned, previous work (Fontal et al., 2021) explored climatic influences on COVID-19 spread, revealing substantial negative correlations between temperature, absolute humidity, and the initial growth rate of COVID-19 cases globally. Highlighted transient negative associations between COVID-19 cases and temperature across different regions and seasons, with specific temperature (12 − 18^°^*C*) and absolute humidity (4 − 12*g/m*^3^) ranges were described as intervals where transmission rates in the population were heightened.

In this study, we investigate the relationship between climatological variables—particularly temperature and absolute humidity—and the incidence of the SARS-CoV-2 Omicron variant (*B*.1.1.529), with a focus on their potential role in shaping transmission dynamics. Leveraging the Scale-Dependent Correlation (SDC) methodology, we examine how these associations vary across spatial and temporal scales, identifying the climatic ranges where significant correlations with COVID-19 incidence are most frequently observed. This approach enables us to track the transient and scale-specific nature of the climate-epidemic relationship across global and regional contexts.

To complement the statistical analysis, we implement a suite of stochastic, population-based compartmental models under different assumptions for the infection rate (constant, seasonal, and climate-driven). By comparing model performance, we assess the added value of incorporating climatic covariates—particularly temperature—in explaining the dynamics of Omicron spread.

Through this dual framework—SDC and dynamical modeling—we aim to: (i) characterize the climate sensitivity of Omicron incidence, (ii) evaluate the consistency of such sensitivity across different geographic regions, and (iii) examine its temporal modulation, especially regarding seasonal effects. Ultimately, our results contribute to a more nuanced understanding of climate-mediated epidemic control, with implications for regional risk assessment and preparedness in the context of ongoing and future COVID-19 waves.

This paper is presented as follows:

## 2. Methods

### 2.1. Data

The circulation of strains was obtained from the GISAID hCoV-19 Dataset. GISAID, a global scientific initiative established in 2008, provides open access to genomic data for influenza viruses and SARS-CoV-2. It has been instrumental in the COVID-19 response, offering the first complete SARS-CoV-2 genome sequences on January 10, 2020. GISAID data includes virus genetic sequences, which are crucial for tracking variants like Omicron *B*.1.1.529.

As illustrated in Figure S2, the B.1.1.529 variant,first sequenced in November 2021, rapidly became dominant within a month (starting in the penultimate week of December 2021). Its peak prevalence occurred from January 2022 to November 2022, after which sequencing data indicated a decline in its predominance by the last week of January 2023 (Figure S2a). The timeline marking the onset and subsequent decline of Omicron’s prevalence closely aligns with global trends in daily COVID-19 cases (Figure S2b). This study will specifically examine this critical period. The gray shaded area in the figure denotes the standard deviation around the global mean of daily new cases per 100,000 individuals, providing insights into the variability among countries during the specified timeframe.

ERA5, the fifth generation of ECMWF atmospheric reanalysis, provides comprehensive global climate data from 1940 to the present through the Copernicus Climate Change Service (C3S) API. For this study, data for 2m Dewpoint Temperature, 2*m* Temperature, Mean Sea Level Pressure, and Total Precipitation were extracted and processed into daily averages. Additionally, Relative Humidity and Absolute Humidity were derived, offering insights into climate impacts on virus capacity of transmission.

NASA’s Socioeconomic Data and Applications Center (SEDAC) provides the Gridded Population of the World (GPW), version 4, which offers high-resolution global population density data, including for the year 2020. This data is used to compute weighted averages of meteorological variables per regional unit based on population density, ensuring accurate reflection of the impact on COVID-19 transmission at the population level.

Our World in Data (OWID) provides detailed data on COVID-19, including confirmed cases, deaths, tests, and vaccinations, sourced from the WHO COVID-19 Dashboard. Data preprocessing involves generating a complete daily series by assuming an equal distribution of weekly reported cases and calculating a centered 7-day rolling average to smooth out daily variations, as the study addresses weekly-scale variations.

### 2.2. Statistics and mathematical methods

#### 2.2.1. Scale-Dependent Correlation (SDC) Analysis

To identify transient associations between climatological variables and COVID-19 incidence, we used Scale-Dependent Correlation (SDC) analysis Rodó and Rodríguez-Arias (2006). SDC is a time series method designed to detect localized, short-lived correlations at varying temporal scales. This allows us to uncover transient relationships that may be hidden in full-series or static correlation metrics.

The analysis involves iteratively extracting fragments of fixed length *s* from two time series (e.g., temperature and incidence), and computing their correlation using Pearson’s or Spearman’s method. A randomization test with *m* permutations is used to determine the statistical significance of observed correlations. Results are displayed as time–scale correlation maps (SDC plots), where significant associations are visualized across different time windows and fragment sizes.

Key methodological parameters such as fragment length, correlation type, and the treatment of significance thresholds were calibrated based on sensitivity analyses. Full algorithmic details and mathematical formulations are provided in the Supplementary Material.

#### 2.2.2. Population-Based Model

To complement the statistical analysis, we implemented a population-based stochastic compartmental model to simulate COVID-19 dynamics under the influence of climatological variables. The model builds upon a modified SEIR framework, extending it into an SEIQRDPV structure to incorporate vaccination stages, protected populations, and a temperature-modulated infection rate.

The infection rate is modeled as a time- and temperature-dependent function, allowing for seasonal and climatic effects to influence disease spread.

This formulation enables the evaluation of how fluctuations in ambient temperature modulate the force of infection over time. Model compartments include vaccinated individuals at various dose levels, detected and undetected infections, recovered, deceased, and protected populations.

For full details on the compartment definitions, transition equations, and parameterization, refer to the Supplementary Material.

## 3. Results

### 3.1. Scale-Dependent Correlation (SDC) Analysis

#### 3.1.1. Global Scale Analysis

To find out the potential relationship between climate variables (specifically here temperature *T* and absolute humidity *AH*) and COVID-19 incidence during the predominance of the Omicron *B*.1.1.529 variant, a two-way SDC analysis was conducted. This analysis considered daily new COVID-19 cases across different regions from December 19*th*, 2021 to January 31*st*, 2023.

The SDC method utilized a temporal window size *s* of 105 days, with varying lags between the placement of windows in the COVID-19 cases and climate variable time series (from 0 to 21 days). The resulting plots in Fig. 4 display scale-dependent correlations between these variables across different geographic regions, grouped by latitude and longitude.

In the SDC plots, the left side verticla plot displays the time series of daily new COVID-19 cases (time goes from top to bottom). This time series illustrates the daily fluctuations in COVID-19 incidence over the study period, providing a clear view of how the number of cases changed over time.

The top series of the plots shows the time series of the climate variable (time going left to right), which can be either temperature (*T*, for Fig. 4) or absolute humidity (*AH*, in Fig. 5). Similar to the COVID-19 cases, this time series presents the daily variations in the selected climate variable, allowing us to observe how climatic conditions change over time.

In the central grid of the plot, each square cell is colored according to the Spearman correlation coefficient. This coefficient measures the strength and direction of the relationship between COVID-19 cases and the climate variable (red for negative correlations and blue for positive ones). The rows and columns of this grid correspond to the positions of the two respective time windows, each of size *s*, along the time series. The distance from the diagonal of the grid represents the lag between the positions of these windows, ranging from 0 to 21 days. For example, a cell exactly on the diagonal indicates no lag between the windows, while a cell of the diagonal indicates that one time series is leading or lagging the other by a certain number of days.

The panel below each plot displays the maximum correlation coefficient obtained at each time point (vertically, relative to the time of the climate time series). Only correlations found to be significant in a non-parametric randomized test (*α* = 0.05) are shown and colored, with red (blue) points corresponding to negative (positive) correlations.

To illustrate this further, consider an example: if a significant blue point appears in the central grid when the COVID-19 window is in March and the climate window is in February (indicating a 30-day lag), this suggests a positive correlation between the climate variable in February and the COVID-19 cases occurring in March. This delay might imply that specific climatic conditions in February are associated with an increase in COVID-19 cases one month later, a timing that should involve all the cascading effects contained in the epidemiological cycle (e.g. becoming infected and developing infectiousness, contagion and appearance of infected persons in hospitals).

Significant findings in Fig. 4 and 5 include strong transient negative correlations observed in Northern Hemisphere regions during the initial rise of Omicron in late 2021 and up until May 2022, right at the midway of Spring. Transient, not consistent and weaker, positive correlations are also detected during the warmer months (June 2022), expressed as a break for some locations, from which negative associations are shown again for some locations during July-September 2022. It should be noted that this method does not imply causation, but indicates potential intervals of true association, meaning some others might be spurious or arising from coincidental dynamics in the evolution of time series. Concommitant alternative approaches should then be employed to further confirm or deny the presence of real mechanistic associations between the tested variables (this will be done in this study with the use of an epidemiological model in the sections below).

For Southern Hemisphere regions, strong transient negative correlations were observed from May 2022 to September 2022 (i.e. coinciding also with colder months in the Southern Hemisphere),flanked by positive correlations before and after these periods.

Similar findings were found with absolute humidity *AH*, which is expected due to the large dependence and correlation between these two humidity variables.

#### 3.1.2. Country-Specific Analysis

Further analyses were conducted at the country scale, focusing in this section on European Union (EU) countries.

In the plots displayed, panels from SDC analysis for the maximum absolute correlations between daily time series for the specific meteorological variable (*AH*, left; *T*, right) and COVID-19 new cases, as a function of time. These panels correspond to those shown below the SDC grid of local correlations in Fig. 1 and 2, but here at the country level for five European countries (Germany, France, Italy, Spain, and the United Kingdom). As before, *s* = 105, and the maximum correlation is chosen for lags from 0 to 21 days in accordance with incubation times. Colors correspond to maximum negative (red) and positive (blue) correlations for each day.

**Figure 1.**
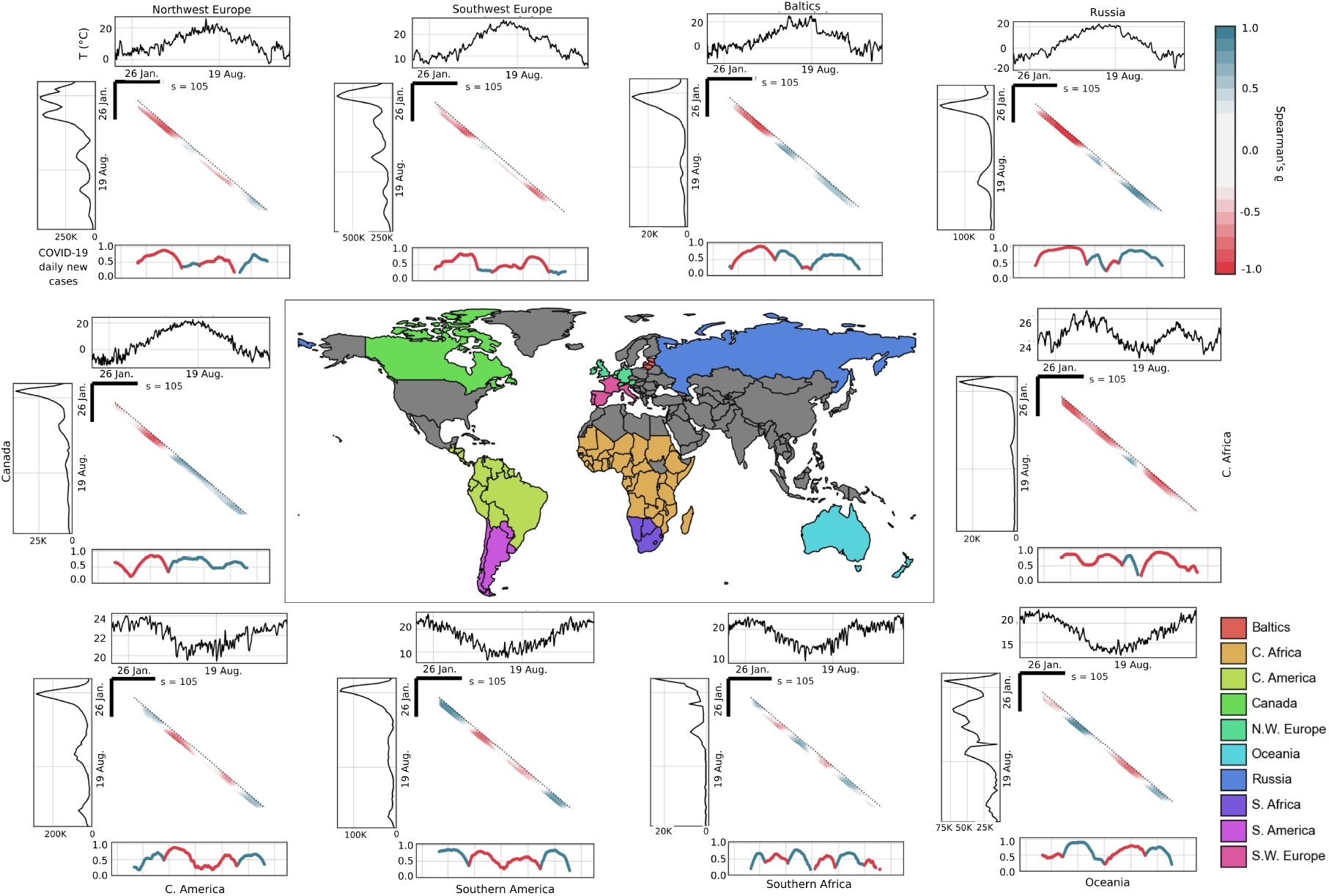
SDC analyses for COVID-19 new daily cases and temperature for grouped countries

**Figure 2.**
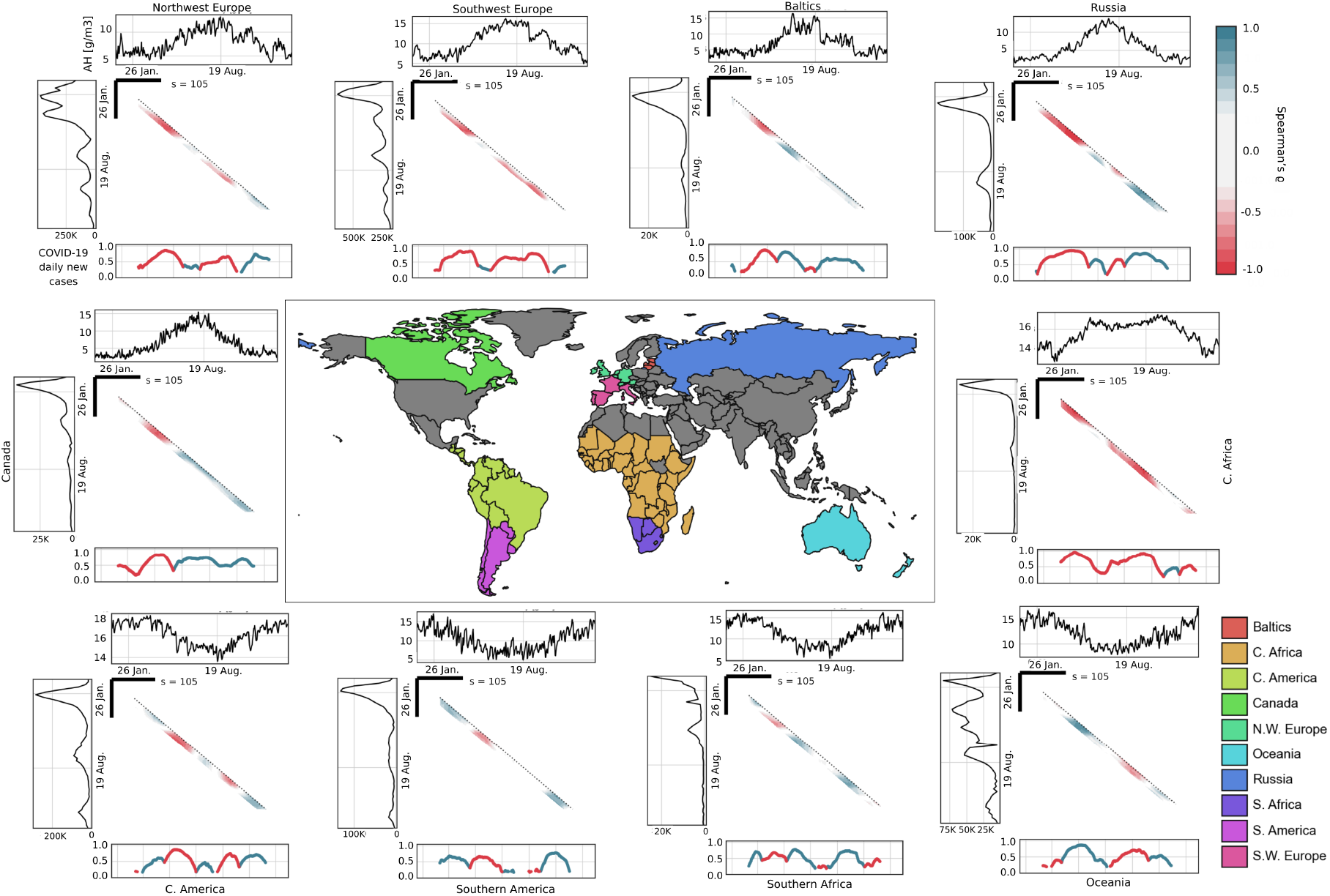
SDC analyses for COVID-19 new daily cases and absolute humidity for grouped countries

Negative relationships occur largely in synchrony across different countries, during the same time intervals: for example, during the rise of the Omicron wave in the fall as *T* and *AH* fall, and during the waning of the same wave as *T* and *AH* rise, with a break in between (Figure 3).

**Figure 3.**
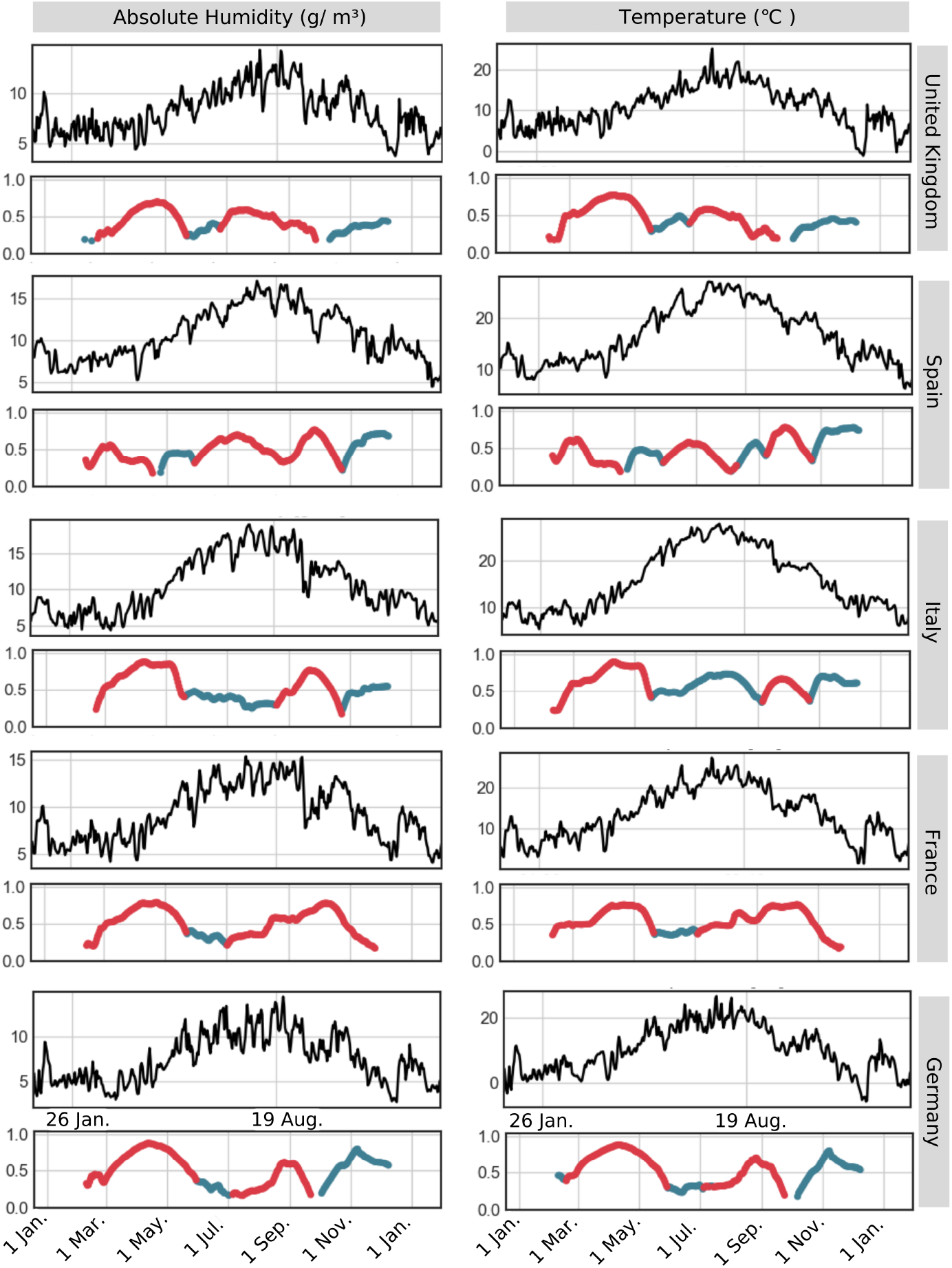
Transitory correlations between meteorological factors and COVID-19 incidence for EU countries

**Figure 4.**
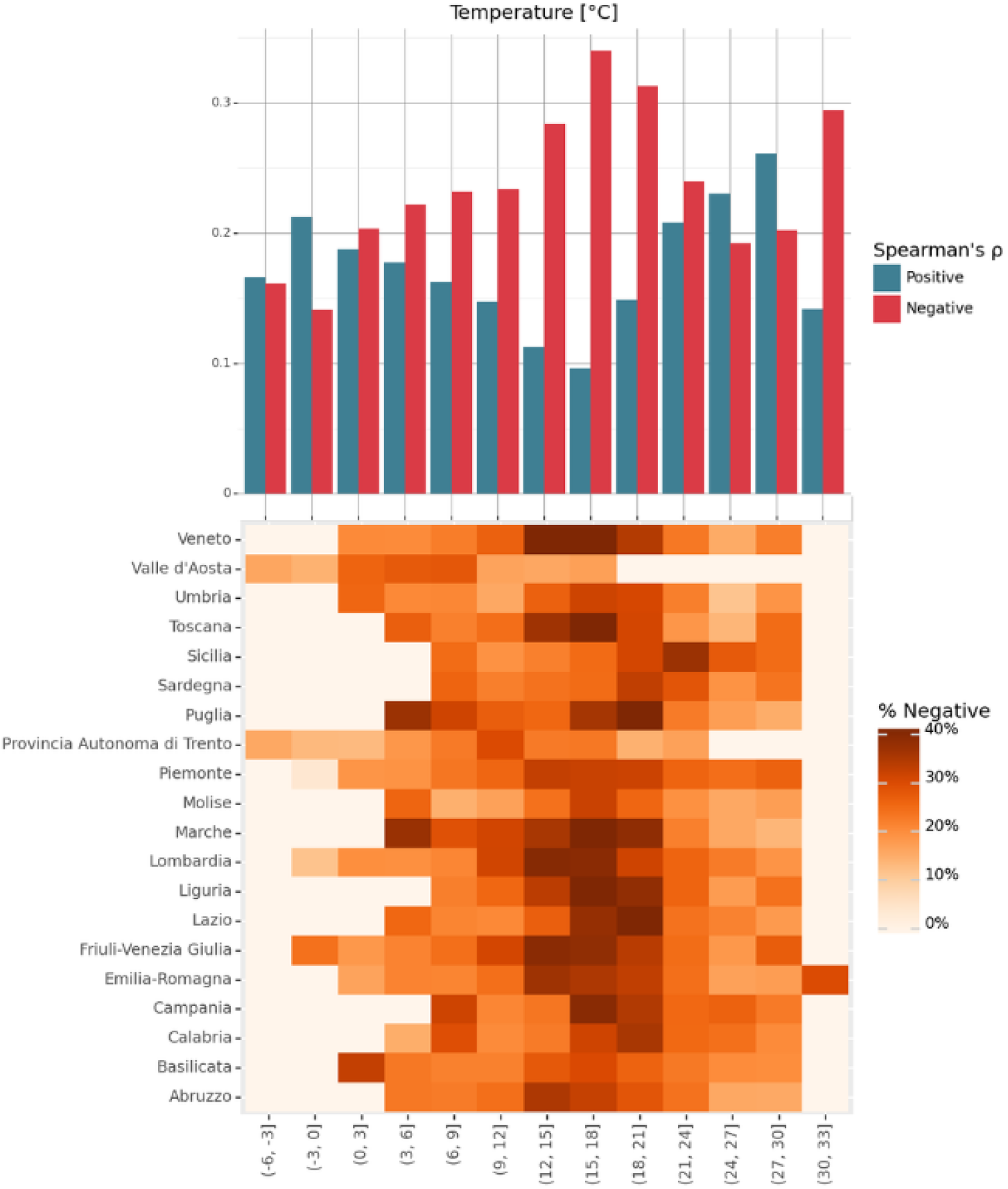
Distribution of correlations between COVID-19 cases and temperature (*T*) across Italian regions

**Figure 5.**
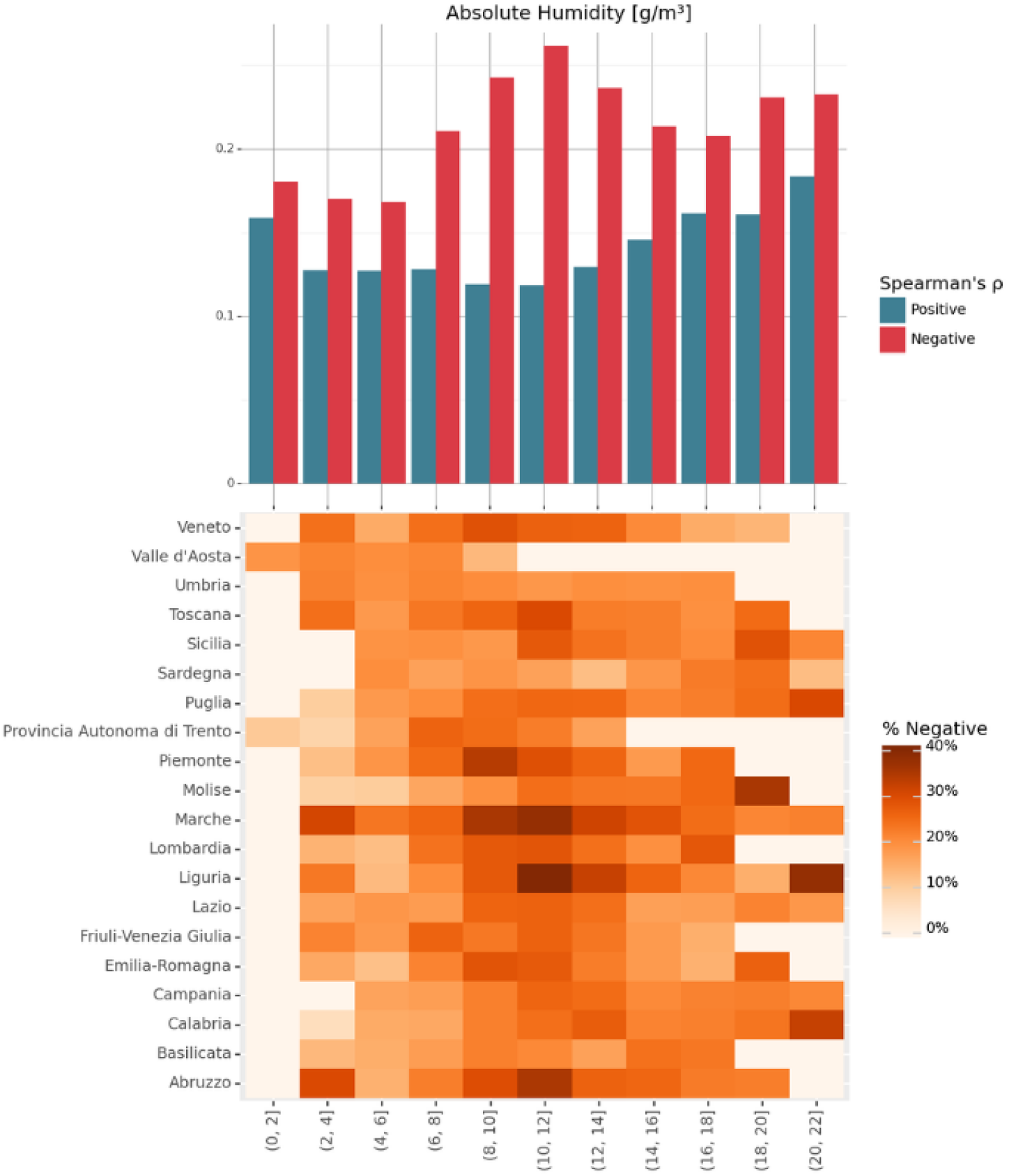
Distribution of correlations between COVID-19 cases and absolute humidity (*AH*) across Italian regions

Next, it was of interest to investigate if, as formerly shown for the first COVID-19 variants, potential threshold values at the population level could be identified in both temperature (*T*) and absolute humidity (*AH*), that might be associated with the ulterior spread of COVID-19. This analysis aimed to identify specific ranges of these climate variables that exhibit non-stochastic correlations with COVID-19 cases, indicating ranges for heightened effects, and to determine whether correlations diminish outside these ranges.

The analysis utilized data from all regions of Italy, leveraging detailed regional COVID-19 case data. Correlations between COVID-19 cases and each climate covariate (*T* or *AH*) were collectively analyzed across Italian regions using a window size *s* of two weeks (14 days).

The resulting distributions (Figures 4 and 5) illustrate the proportion of all possible correlations (pairs of time intervals between the two time series) and their excess (negative to positive or vice versa) falling within specific ranges of each specific climate covariate. These distributions categorize correlations into significant positive and negative associations and include also non-significant correlations.

The peaks in these distributions again clearly indicate intervals of temperature and absolute humidity where the effects of climate on COVID-19 transmission are most pronounced. Specifically, these peaks highlight thresholds in temperature and absolute humidity ranges that show the strongest correlations with COVID-19 case numbers. For temperature, the critical range is found to be between 12−21 °*C* . This suggests that within this temperature range, there is a more significant relationship between the temperature and the number of COVID-19 cases and that environmental conditions appeared more suited for the spatial and temporal propagation of the virus in the population. Similarly, for absolute humidity, the range of 8–12 g/m^3^ has been identified as particularly influential. This means that when absolute humidity falls within this range, the likelihood of COVID-19 transmission at the population level is more facilitated. These findings help to pinpoint specific climate conditions under which COVID-19 transmission dynamics are affected the most, thereby providing valuable insights for public health strategies and interventions. A noticeable degree of coincidence between earlier variants of the virus and the Omicron underscore how similar meteorological conditions seem to affect the different strains, despite the inter-strain differences that may exist.

### 3.2. Population-based model fitting and performance

The model was able to reproduce observed epidemic trends under different climatic regimes. A full description of the model’s structure and parameter calibration is provided in the Supplementary Material. This model was implemented using Python and calibrated with a least squares algorithm from the SciPy library. The error function minimized by this algorithm was derived from the normalized residuals of total cases (*Q* + *R* + *D*), deaths (*D*), and the vaccinated population (*V*_*i*_, where *i* = 1,2,3).

Figures 6, 7 and 8 present the model performance under three infection rate assumptions: constant, temperature-based, and seasonal. The first two panels in each figure correspond to global values: the top panel shows the global mean of active cases, and the middle panel shows the corresponding global normalized error. The bottom panel breaks down model performance by continent, allowing us to assess regional variation in fit quality.

**Figure 6.**
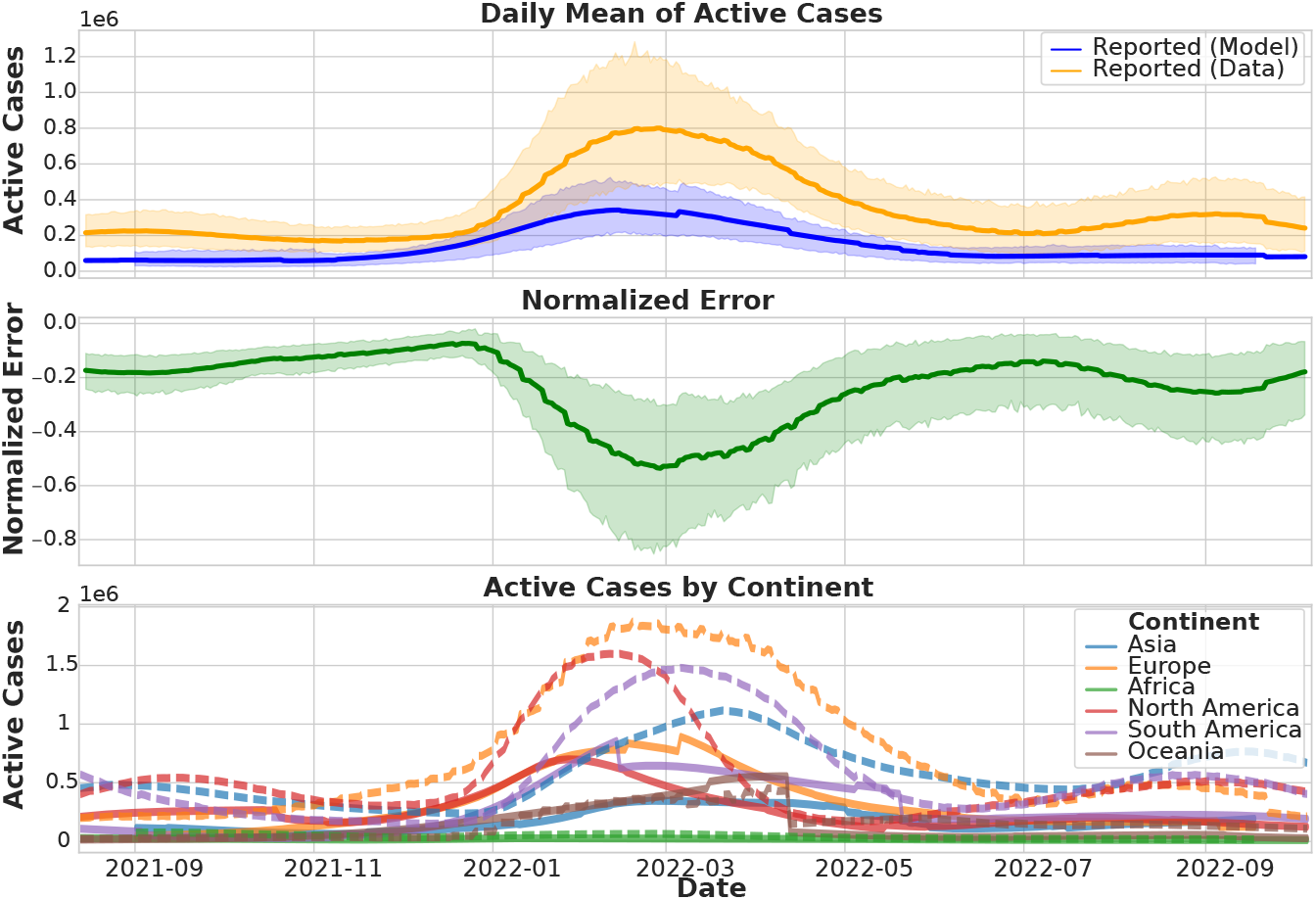
Comparison of the model’s predictions of active COVID-19 cases using a constant infection rate, with real reported data over a subperiod focused on the main Omicron wave. The top panel shows the global daily mean of active cases (model vs. reported). The middle panel presents the global normalized error over time. The bottom panel displays active cases by continent, with solid lines for model predictions and dashed lines for reported data. This allows assessment of performance and discrepancies across different regions.

**Figure 7.**
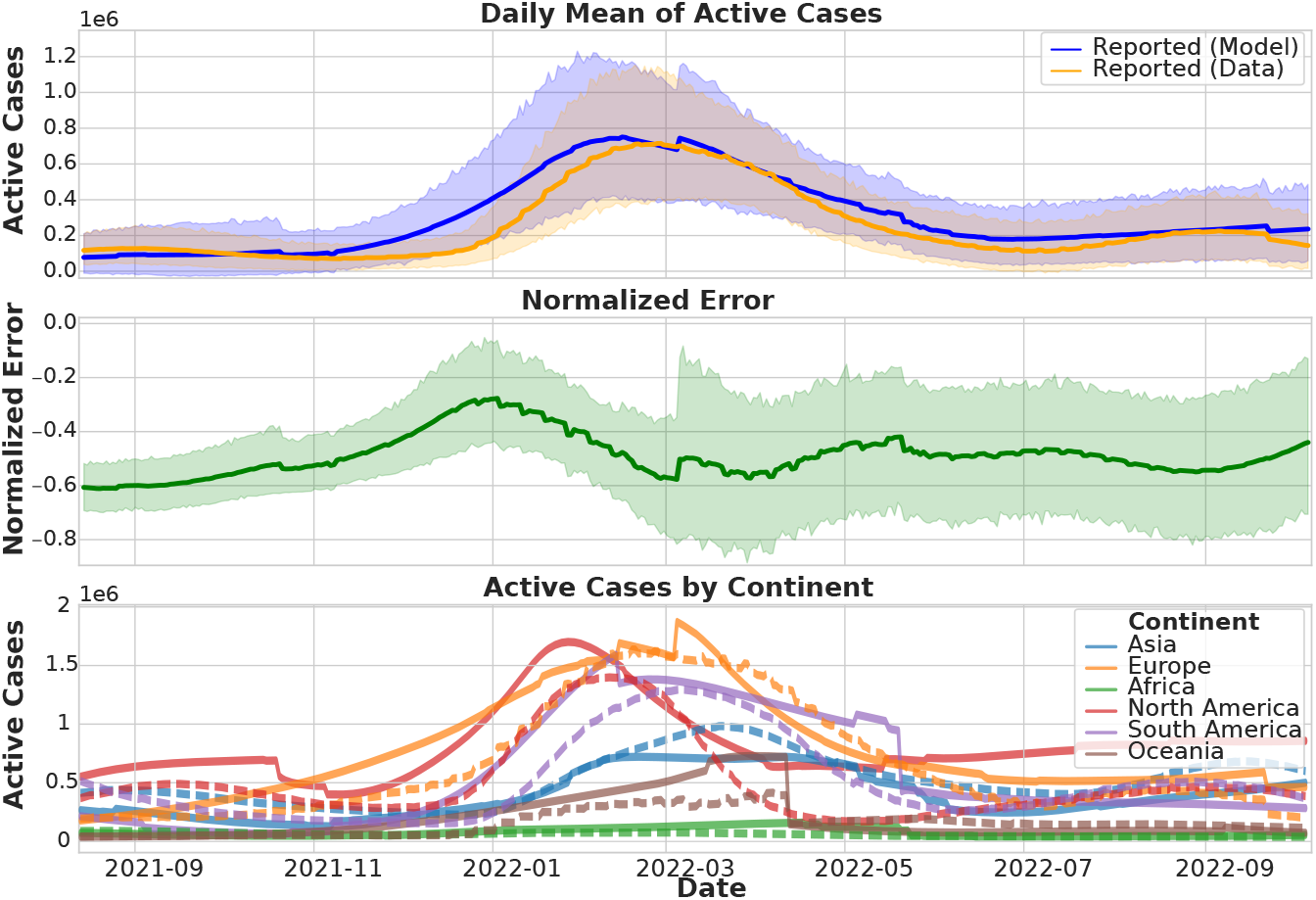
As in Fig. 6, but using an infection rate modulated by temperature variations, with real reported data over a subperiod focused on the main Omicron wave. The top panel shows the global mean of active cases; the middle panel illustrates global normalized errors. The bottom panel provides active cases by continent, highlighting the improved predictive accuracy of the temperature-driven model and regional variability in modelfit.

**Figure 8.**
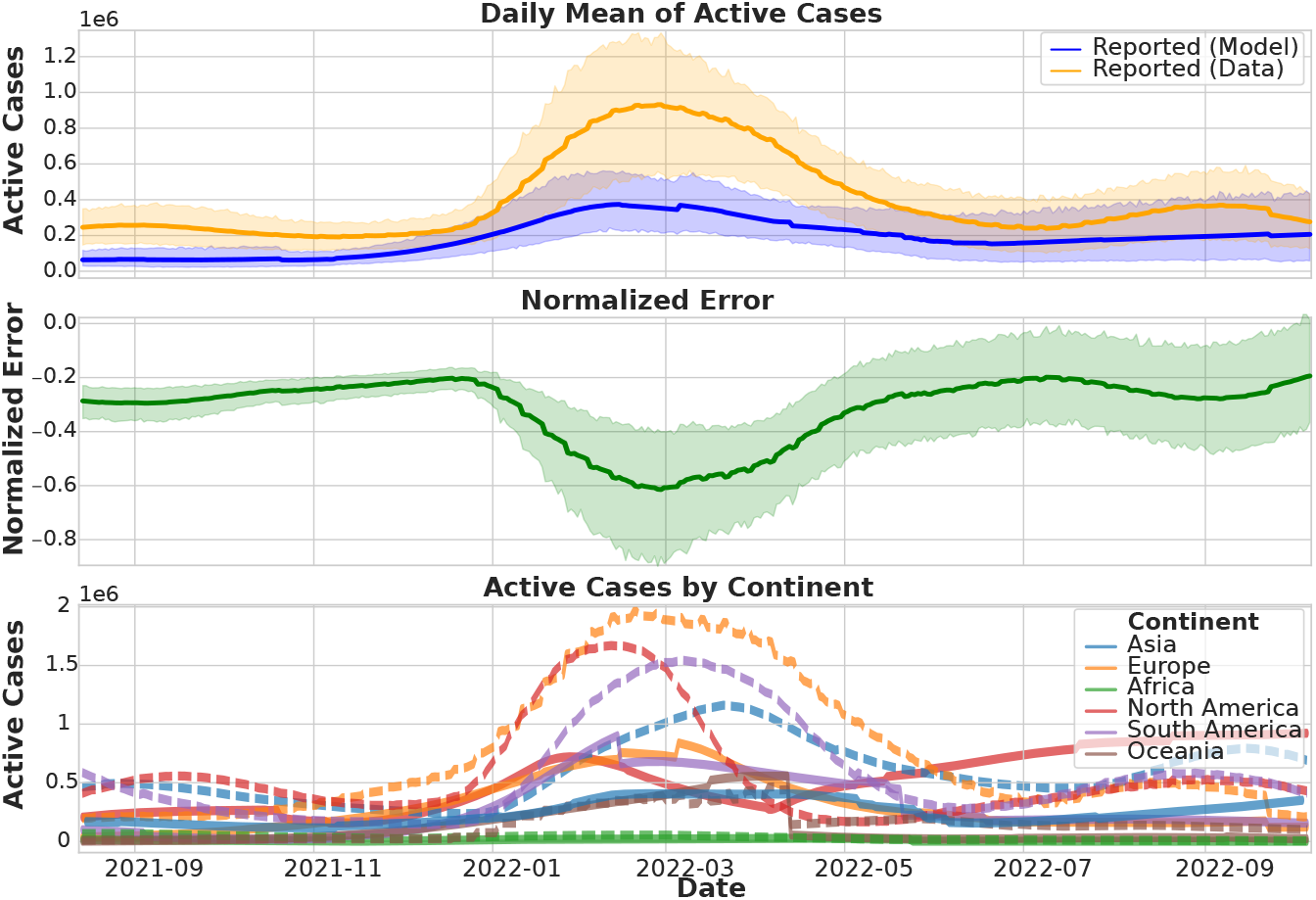
As in Fig. 6, but using a seasonal infection rate, with real reported data over a subperiod focused on the main Omicron wave. The top and middle panels represent global-level predictions and errors, respectively. The bottom panel shows continent-level comparisons, illustrating how seasonal dynamics influence model accuracy across regions.

The model with temperature dependency (Figure 7), which incorporates temperature as a factor influencing the infection rate, emerges as the most accurate approximation among the three models. This model not only captures the global dynamics more effectively but also demonstrates superior accuracy at the continental level. The improved performance is evident in the top panel of the figure, where the model’s predicted active cases align more closely with the actual reported data compared to the other models. The bottom panel further underscores this accuracy, showing a better fit for active cases by continent.

Although the model including absolute humidity (AH) also yielded good results (not shown), the temperature-based (T) model provided a notably better fit, with lower residual errors and improved stability. This suggests that while both climate variables are relevant, temperature may exert a more consistent and dominant influence in explaining population-level transmission patterns during the Omicron wave.

Moreover, the middle panel, which displays the normalized error, highlights the significant reduction in prediction error achieved by the temperature approach. By focusing on the central portion of the error graph, it becomes apparent that the temperature-based model reduces the error by approximately 75%compared to the other models.

## 4. Discussion and Conclusions

The investigation into the relationship between climate variables, specifically temperature (*T*) and absolute humidity (*AH*), and COVID-19 incidence during the predominance of the Omicron *B*.1.1.529 variant reveals significant correlations and highlights the same role described to be exerted by physical conditions on the spread and viability of viruses attached to aerosol particles as potentially crucial factors. The analysis conducted from December 19*th*, 2021, to January 31*st*, 2023 as well as the fitted COVID-19 models, indicates that climatic factors played again a crucial role in modulating COVID-19 transmission at the population level.

In the Northern Hemisphere, strong transient negative correlations were observed during the initial rise of Omicron in late 2021, while positive correlations were detected during the warmer months. This pattern indicates a complex interaction between climate and virus transmission dynamics. The synchrony in negative correlations across different European Union countries during the rise and fall of the Omicron wave reinforces the consistency of climatic influence on COVID-19 spread across various spatial and temporal scales.

Furthermore, identifying specific thresholds for temperature and absolute humidity, where transmission effects are most pronounced, emphasizes the importance of these climatic parameters. These findings suggest that public health strategies should consider meteorological conditions, especially by improving indoor ventilation and adapting control measures according to climate variations.

Alongside, the application of the *SEIQRDPV* model, which incorporates temperature as an infection rate modulator, effectively reproduced the real dynamics worldwide, even with the large heterogeneity observed in the data sets. The model demonstrated considerable stability within the fitted infection rate values, validating its robustness and reliability. This stability underscores the model’s capability to accurately predict infection trends, even under varying climatic conditions and data uncertainties.

Overall, these results support the existence of strong external drivers of COVID-19 transmission, similar to the seasonality observed in influenza and other coronaviruses. The significant correlations between temperature, absolute humidity, and COVID-19 incidence underscore the transient but influential role of climate in virus spread. These findings align with previous studies, despite methodological differences, and highlight the need for incorporating meteorological parameters into public health planning.

Future public health prevention strategies could highly benefit from tailored climate services and early-warning systems to better manage waves of COVID-19 transmission, especially as new variants with higher transmissibility and immune evasion capabilities emerge. The persistence of climatic influence on COVID-19 transmission irrespective of the variant under investigation, further reinforces the necessity for ongoing public health interventions to include meteorological indicators as a promising way to anticipate impacts and help mitigate the impact of the virus effectively. These insights emphasize that public health strategies must remain adaptive and incorporate climatic factors to respond to the evolving challenges posed by COVID-19 new variants and potentially also to the threats of new respiratory viruses.

## 5. Acknowledges

## 6. Code and data Dvailability

